# SenLect: a genetically encoded system to purify senescent cells

**DOI:** 10.1101/2025.09.09.675259

**Authors:** Hourieh Tousian, Rachael A Protzman, Timothy J. Sargeant, Julian M. Carosi

## Abstract

Cellular senescence is a state of irreversible cell cycle arrest that prevents cancer and promotes biological ageing and inflammation. The molecular landscape of cellular senescence has been met with conflicting findings, likely due to contamination of senescent cells by neighbouring proliferating cells. Here, we developed a genetically encoded and tractable system to purify senescent cells in culture termed ‘SenLect’ (<u>Sen</u>escence se<u>Lect</u>ion). SenLect is a genetic cassette that expresses mCherry-2A-PuroR under the control of the miR-146a senescence-activated promoter, which provides senescent cells with a selective advantage over proliferating cells in the presence of the antibiotic puromycin. We validated this one-step system in primary HUVECs and HeLa cervical cancer cells to purify live senescent cells following DNA damage (UV irradiation, hydrogen peroxide, and doxorubicin) and cell-cycle inhibition (palbociclib). Puromycin-selected cells were enriched for various senescence-related phenotypes, including cell cycle arrest, increased cell size, and lysosomal β-galactosidase activity. By using SenLect to remove proliferating cells, we uncovered how the proteome is rewired during senescence induced by palbociclib. This improved the detection of protein signatures linked to the senescence-associated secretory phenotype (SASP), and metabolic shifts from mitochondrial respiration to glycolysis, and from nucleotide synthesis to catabolism. It also revealed sub-proteome level rewiring of organelles such as mitochondria, lysosomes and the nucleolus, as well as protein cohorts like kinases and core-essential proteins. SenLect provides a consistent and reliable method for isolating senescent cells that is suitable for downstream applications or analyses.

## INTRODUCTION

Cellular senescence is a permanent pause in the cell cycle that prevents cancer formation and promotes tissue repair ^1^. Senescent cells release inflammatory factors to attract the immune system and promote their clearance. Increased formation or reduced clearance of senescent cells leads to accelerated biological ageing ^1^. So far, there is no single marker for senescent cells ^2-4^. Hence, the evaluation of different markers is required to confirm this state. The most common markers of senescent cells are large cell size, high granularity, lysosomal β-galactosidase activity, and increased levels of cell cycle inhibitor proteins such as p21, p16, and p53 ^4^. There is significant heterogeneity among senescence inducers, with even the same inducer yielding varying proportions of senescent cells across different cell lines ^5-7^. Most studies are therefore conducted on mixed cultures of senescent and proliferative cells ^8-10^. This poses challenges for data interpretation because proliferative cells can contaminate and overtake cultures of senescent cells, which do not divide. This makes it difficult to determine whether treatments actually return senescent cells to a proliferative state, prevent the formation of new senescent cells from proliferative cells, or just kill senescent cells. It also limits the ability to study the molecular landscape of this cell-state transition.

Various methods have been employed to isolate and maintain live senescent cells. The most common approach for isolating senescent cells is fluorescence-activated cell sorting (FACS), which typically uses parameters such as cell size, autofluorescence ^11,12^, and lysosomal β-galactosidase activity ^13^, or a combination thereof ^12,14^. However, FACS can be stressful for cells, especially for senescent cells which are fragile, leading to significant cell loss. Additionally, the parameters used in FACS are not entirely specific to senescence, which may impact the purity of isolated cells. To overcome these limitations, we developed a genetic system capable of directly purifying senescent cells away from proliferative cells in culture. We took advantage of the high miR-146a expression levels found in senescent cells of various tissue origins ^15,16^. We designed a genetically encoded system termed ‘SenLect’ (<u>Sen</u>escence se<u>Lect</u>ion) that expresses mCherry and puromycin resistance under the control of the senescence-activated miR-146a promoter. After senescence induction, proliferative cells are selectively eliminated by the antibiotic puromycin, enabling enrichment of senescent cells for downstream applications.

## RESULTS

### SenLect enriches for the low-frequency senescent cell population that arises under basal conditions within proliferating primary cultures

We developed a genetically encoded promoter-based reporter to enable the identification and selection of senescent cells in culture that we have termed ‘SenLect’. SenLect encodes mCherry-P2A-PuroR (puromycin resistance; puromycin N-acetyltransferase) under the control of the senescence-inducible miR-146a promoter (-1.5 kb from the 5’ region of exon 1) ^15,16^ **(Figure 1A)**. It also encodes TagBFP-T2A-BlastR (blasticidin S resistance; blasticidin S deaminase) that is constitutively expressed by the CMV promoter **(Figure 1A)**. SenLect is packaged within a lentiviral transfer plasmid (i.e., pLV[Exp]) that is used to produce lentivirus, infect target cells, and select for its stable genomic integration with blasticidin S (or TagBFP fluorescence) **(Figure 1B)**. The miR-146a promoter is strongly activated in senescent cells, but is only weakly active or inactive in proliferating cells **(Figure 1A and B)** ^15,16^. Importantly, the miR-146a promoter is activated after cells enter senescence, rather than immediately in response to the triggering event ^15,16^. Therefore, SenLect enables mCherry fluorescence-based detection of senescent cells and their enrichment via puromycin-mediated elimination of proliferative cells **(Figure 1A and B)**. We also generated control plasmids to facilitate accurate gate setting during flow cytometry analysis of the SenLect system **(Figure S1)**. These included a plasmid in which mCherry-P2A-PuroR is constitutively expressed under the UbC promoter **(Figure S1)** and a plasmid lacking a promoter, resulting in no mCherry-P2A-PuroR expression **(Figure S1)**.

**Figure 1.**
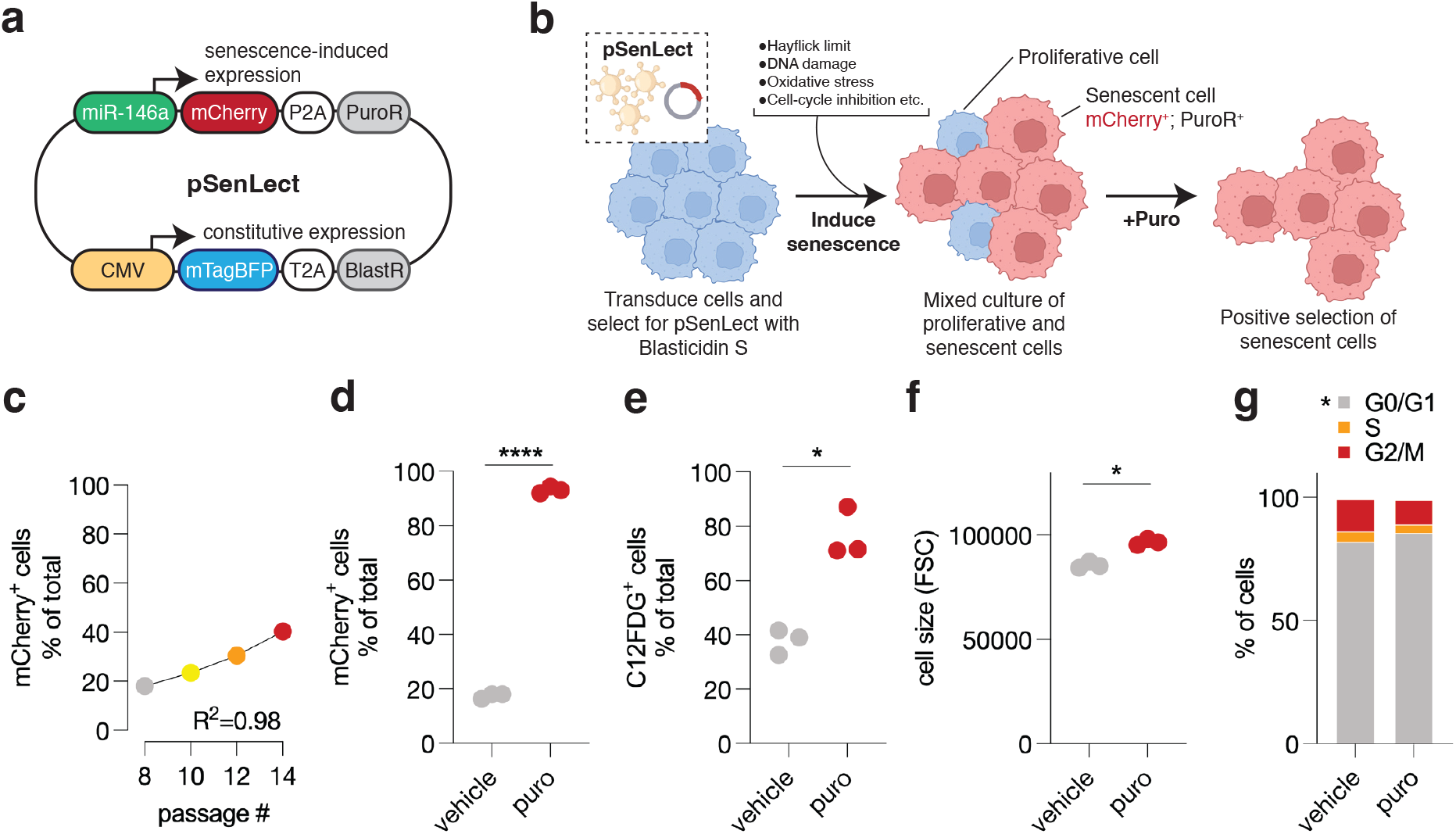
SenLect enriches for the low-frequency senescent cell population that arises under basal conditions within proliferating primary cultures. A. Summary of the SenLect plasmid (pSenLect). B. Summary of how SenLect enables selection of senescent cells via puromycin (puro)-based elimination of proliferative cells. C. Proportion of mCherry^+^ cells with increasing passage number in pSenLect-integrated primary HUVEC cells. n=4 independent experiments. Pearson correlation analysis. D. Proportion of mCherry^+^ cells after selection with or without puro in pSenLect-integrated primary HUVEC cells. n=3 independent experiments. Paired t-test. E. Proportion of C12FDG^+^ cells after selection with or without puro in pSenLect-integrated primary HUVEC cells. n=3 independent experiments. Paired t-test. F. Cell size (FSC) after selection with or without puro in pSenLect-integrated primary HUVEC cells. n=3 independent experiments. Paired t-test. G. Proportion of cells in cell cycle stages after selection with or without puro in pSenLect-integrated primary HUVEC cells. Data represents mean values from n=3 independent experiments. Two-way ANOVA with Sidak’s multiple comparisons test.

To test the SenLect system, we transduced primary Human Umbilical Vein Endothelial Cells (HUVECs) at passage five then selected for integration with blasticidin S. We first sought to establish whether SenLect could identify and enrich for senescent cells under normal growth conditions. As cells aged (i.e., increasing passage number), we observed a linear increase in the proportion of mCherry^+^ senescent cells within the culture, as measured by flow cytometry **(Figure 1C)**. Selection with puromycin increased the proportion of mCherry^+^ senescent cells from 17.5% to 93.1%, doubled the proportion of cells with high senescence-associated β-galactosidase activity as determined by 5-dodecanoylaminofluorescein di-β-D-galactopyranoside (C12FDG) cleavage, and increased the size of surviving cells (forward scatter, FSC) **(Figure D-F)**. A modest increase in the proportion of cells arrested in the G0/G1 phase of the cell cycle was noted after selection **(Figure 1G)**. Thus, SenLect can be used to enrich for the low abundance of senescent cells that exist in proliferating cell cultures under basal conditions.

### SenLect enriches for the low-frequency senescent cell population that forms after exposure to oxidative stress

We next sought to determine how SenLect responds to established inducers of senescence in HUVECs. UV irradiation (312 nm), oxidative stress with H_2_O_2_ and chemotherapy-induced DNA damage with doxorubicin all increased in the proportion of cells in the G2/M phase of the cell cycle **(Figure 2D, H and P)**. Each of these treatments increased the proportion of mCherry^+^ senescent cells, C12FDG^+^ cells, and increased cell size (FSC) **(Figure 2A-C, E-G and M-O)**. In H_2_O_2_-treated cells, these senescence-related phenotypes were enriched after puromycin selection **(Figure 2I-L)**. In contrast, puromycin selection only increased the proportion of mCherry^+^ senescent cells that were treated with doxorubicin, but did not further increase the already high proportion of cells that were C12FDG^+^ or large in size **(Figure 2Q-T)**. Doxorubicin-induced senescent cells accounted for ∼94% of the culture, which may explain why puromycin-based enrichments were ineffective **(Figure 2M-T)**. Thus, SenLect was optimal for selecting small populations of senescent cells within a larger population of proliferative cells.

**Figure 2.**
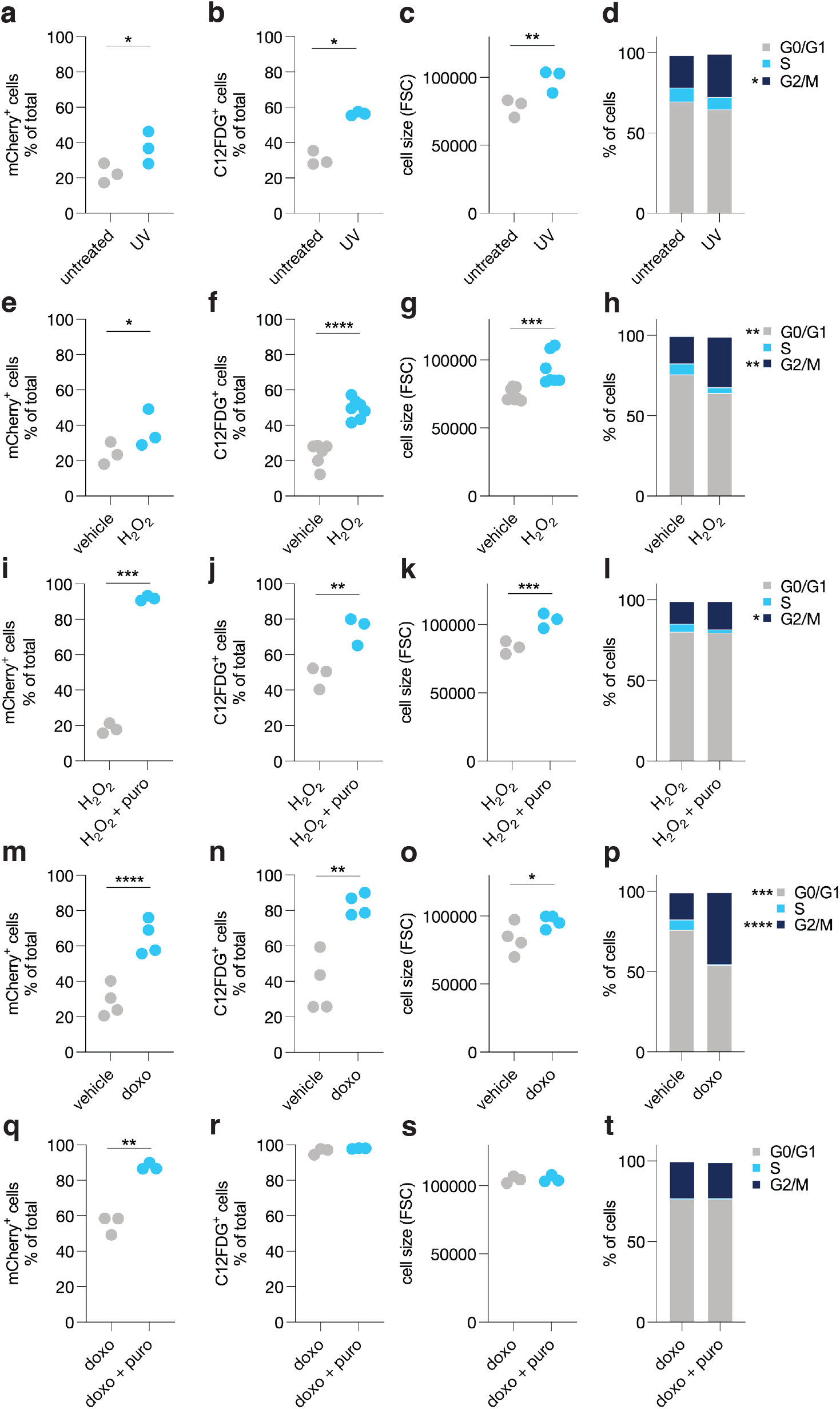
SenLect enriches for the low-frequency senescent cell population that forms after exposure to oxidative stress. A. Proportion of mCherry^+^ cells following treatment with or without UV exposure (312 nm) in pSenLect-integrated primary HUVEC cells. n=3 independent experiments. Paired t-test. B. Proportion of C12FDG^+^ cells following treatment with or without UV exposure (312 nm) in pSenLect-integrated primary HUVEC cells. n=3 independent experiments. Paired t-test. C. Cell size (FSC) following treatment with or without UV exposure (312 nm) in pSenLect-integrated primary HUVEC cells. n=3 independent experiments. Paired t-test. D. Proportion of cells in cell cycle stages following treatment with or without UV exposure (312 nm) in pSenLect-integrated primary HUVEC cells. Data represents mean values from n=3 independent experiments. Two-way ANOVA with Sidak’s multiple comparisons test. E. Proportion of mCherry^+^ cells following treatment with or without H_2_O_2_ exposure in pSenLect-integrated primary HUVEC cells. n=3 independent experiments. Paired t-test. F. Proportion of C12FDG^+^ cells following treatment with or without H_2_O_2_ exposure in pSenLect-integrated primary HUVEC cells. n=7 independent experiments. Paired t-test. G. Cell size (FSC) following treatment with or without H_2_O_2_ exposure in pSenLect-integrated primary HUVEC cells. n=7 independent experiments. Paired t-test. H. Proportion of cells in cell cycle stages following treatment with or without H_2_O_2_ exposure in pSenLect-integrated primary HUVEC cells. Data represents mean values from n=3 independent experiments. Two-way ANOVA with Sidak’s multiple comparisons test. I. Proportion of mCherry^+^ cells following H_2_O_2_ exposure after selection with or without puro in pSenLect-integrated primary HUVEC cells. n=3 independent experiments. Paired t-test. J. Proportion of C12FDG^+^ cells following H_2_O_2_ exposure after selection with or without puro in pSenLect-integrated primary HUVEC cells. n=3 independent experiments. Paired t-test. K. Cell size (FSC) following H_2_O_2_ exposure after selection with or without puro in pSenLect-integrated primary HUVEC cells. n=3 independent experiments. Paired t-test. L. Proportion of cells in cell cycle stages following H_2_O_2_ exposure after selection with or without puro in pSenLect-integrated primary HUVEC cells. Data represents mean values from n=3 independent experiments. Two-way ANOVA with Sidak’s multiple comparisons test. M. Proportion of mCherry^+^ cells following treatment with or without doxorubicin (doxo) in pSenLect-integrated primary HUVEC cells. n=4 independent experiments. Paired t-test. N. Proportion of C12FDG^+^ cells following treatment with or without doxo in pSenLect-integrated primary HUVEC cells. n=4 independent experiments. Paired t-test. O. Cell size (FSC) following treatment with or without doxo in pSenLect-integrated primary HUVEC cells. n=4 independent experiments. Paired t-test. P. Proportion of cells in cell cycle stages following treatment with or without doxo in pSenLect-integrated primary HUVEC cells. Data represents mean values from n=4 independent experiments. Two-way ANOVA with Sidak’s multiple comparisons test. Q. Proportion of mCherry^+^ cells following treatment with doxo after selection with or without puro in pSenLect-integrated primary HUVEC cells. n=3 independent experiments. Paired t-test. R. Proportion of C12FDG^+^ cells following treatment with doxo after selection with or without puro in pSenLect-integrated primary HUVEC cells. n=3 independent experiments. Paired t-test. S. Cell size (FSC) following treatment with doxo after selection with or without puro in pSenLect-integrated primary HUVEC cells. n=3 independent experiments. Paired t-test. T. Proportion of cells in cell cycle stages following treatment with doxo after selection with or without puro in pSenLect-integrated primary HUVEC cells. Data represents mean values from n=3 independent experiments. Two-way ANOVA with Sidak’s multiple comparisons test.

### SenLect enriches for the low-frequency senescent cell population that forms after exposure to palbociclib

Next, we introduced the SenLect system to HeLa cells, an immortal cervical cancer cell line, to overcome technical limitations that arise from replicative senescence. In HeLa cells, doxorubicin resulted in near complete conversion to mCherry^+^ senescent cells, increased the proportion C12FDG^+^ cells, increased cell size (FSC), and increased the proportion of cells in the G2/M phase of the cell cycle **(Figure 3A-D)**. In contrast to HUVECs, neither H_2_O_2_ nor UV irradiation induced senescence in HeLa cells (data not shown). Therefore, we focused on palbociclib, a selective inhibitor of cyclin-dependent kinase 4/6 that has been reported to induce senescence ^17,18^. We arrived at a “goldilocks” dose of palbociclib (7.5 μm) for inducing senescence: low doses failed to induce senescence, and high doses were lethal (data not shown). In HeLa cells, palbociclib led to a minor increase in the proportion of mCherry^+^ senescent cells and increased cell granularity (side scatter, SSC) but did not affect cell size (FSC) or the proportion of cells in the G2/M phase of the cell cycle after a 72-hour treatment window **(Figure 3E-H)**. In contrast, puromycin selection increased the proportion of mCherry^+^ senescent cells that were treated with palbociclib but did not increase cell size **(Figure 3I and J)**. Puromycin selection increased granularity of cells treated with palbociclib compared to those treated with vehicle **(Figure 3K)**. Compared to directly after palbociclib exposure **(Figure 3H)**, following a washout period (without puromycin) there was a reduction in the proportion of cells in the G0/G1 phase of the cell cycle, but an increase in the G2/M phase **(Figure 3L)**. Puromycin selection in palbociclib treated cells increased the proportion of cells in the G2/M phase of the cell cycle **(Figure 3L)**. Thus, SenLect was broadly useful for enriching cell populations with senescent-like properties.

**Figure 3.**
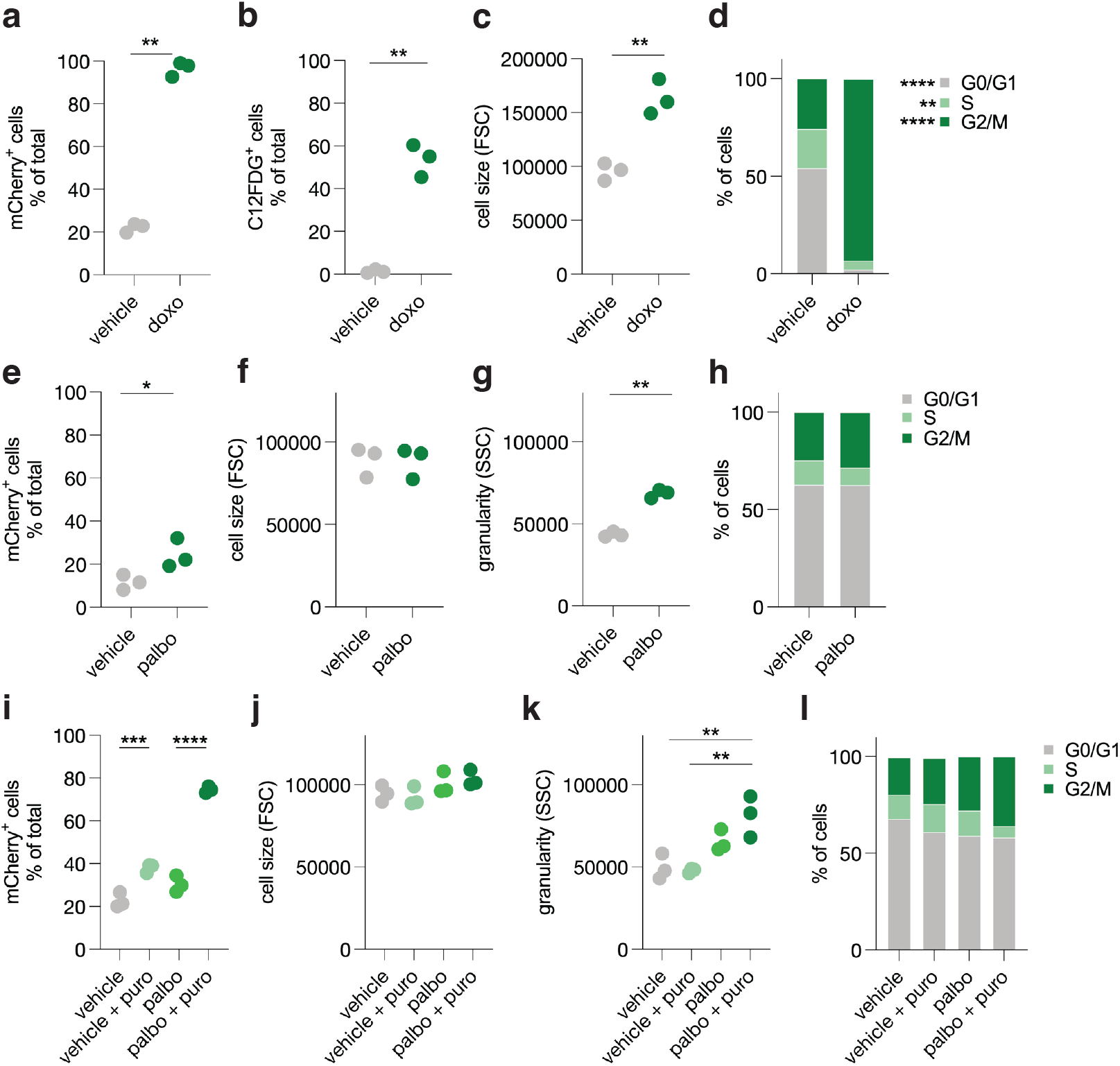
SenLect enriches for the low-frequency senescent cell population that forms after exposure to palbociclib. A. Proportion of mCherry^+^ cells following treatment with or without doxo in pSenLect-integrated HeLa cells. n=3 independent experiments. Paired t-test. B. Proportion of C12FDG^+^ cells following treatment with or without doxo in pSenLect-integrated HeLa cells. n=3 independent experiments. Paired t-test. C. Cell size (FSC) following treatment with or without doxo in pSenLect-integrated HeLa cells. n=3 independent experiments. Paired t-test. D. Proportion of cells in cell cycle stages following treatment with or without doxo in pSenLect-integrated HeLa cells. Data represents mean values from n=3 independent experiments. Two-way ANOVA with Sidak’s multiple comparisons test. E. Proportion of mCherry^+^ cells following treatment with or without palbociclib (palbo) in pSenLect-integrated HeLa cells. n=3 independent experiments. Paired t-test. F. Cell size (FSC) following treatment with or without palbo in pSenLect-integrated HeLa cells. n=3 independent experiments. Paired t-test. G. Cell granularity (SSC) following treatment with or without palbo in pSenLect-integrated HeLa cells. n=3 independent experiments. Paired t-test. H. Proportion of cells in cell cycle stages following treatment with or without palbo in pSenLect-integrated HeLa cells. Data represents mean values from n=3 independent experiments. Two-way ANOVA with Sidak’s multiple comparisons test. I. Proportion of mCherry^+^ cells following treatment with or without palbo after selection with or without puro in pSenLect-integrated HeLa cells. n=3 independent experiments. One-way ANOVA with Sidak’s multiple comparisons test J. Cell size (FSC) following treatment with or without palbo after selection with or without puro in pSenLect-integrated HeLa cells. n=3 independent experiments. One-way ANOVA with Sidak’s multiple comparisons test. K. Cell granularity (SSC) following treatment with or without palbo after selection with or without puro in pSenLect-integrated HeLa cells. n=3 independent experiments. One-way ANOVA with Sidak’s multiple comparisons test. L. Proportion of cells in cell cycle stages following treatment with or without palbo after selection with or without puro in pSenLect-integrated HeLa cells. Data represents mean values from n=3 independent experiments. Two-way ANOVA with Sidak’s multiple comparisons test. G0/G1: vehicle vs. vehicle + puro = ^*^; vehicle vs. palbo = ^**^; vehicle vs. palbo + puro = ^**^; S: vehicle vs. palbo + puro = ^*^; vehicle + puro vs. palbo + puro = ^**^; palbo vs. palbo + puro = ^*^; G2/M: vehicle vs. palbo = ^**^; vehicle vs. palbo + puro = ^****^; vehicle + puro vs. palbo + puro = ^****^; palbo vs. palbo + puro = ^**^.

### Coupling SenLect to global proteomics improves the resolution of senescence-related phenotypes after exposure to palbociclib

We reasoned that by eliminating proliferative cells, SenLect could provide deeper resolution to show how the cellular proteome is rewired during senescence. To monitor this, HeLa cells were treated with or without palbociclib, followed by selection with or without puromycin prior to analysis by whole-cell proteomics. Puromycin-based selection in palbociclib-treated cells led to pronounced proteome remodelling, as indicated by principal component analysis **(Figure 4A)**. To identify which cellular pathways were reprogrammed during senescence, we performed pathway enrichment analysis on differentially expressed proteins (P<0.05; fold change ≥2) using a combination of GO Biological Process Complete ^19,20^ and Reactome ^21^ analyses **(Figure 4B and Table S1)**. For instance, in palbociclib-treated cells after selection, mitochondrial respiration and purine biosynthesis pathways were down-regulated, whereas glycolysis and purine/pyrimidine catabolic processes were upregulated, consistent with known senescence-related metabolic rewiring ^22^ **(Figure 4B and Table S1)**. Gratifyingly, the Reactome pathway “drug-mediated inhibition of CDK4/6 activity” was only enriched in palbociclib-treated cells after selection **(Table S1)**. Examination of organelle-specific sub-proteomes showed that, overall, mitochondrial and nucleolar protein inventories were reduced in palbociclib-treated cells after selection, while lysosomal proteins increased **(Figure S2A)**. Similarly, protein cohorts corresponding to core essential proteins and kinases decreased after selection **(Figure S2B)**. We identified 795 differentially expressed proteins (P<0.05) when comparing palbociclib with vehicle **(Figure 4C and S3, and Table S2)**, which increased to 1660 proteins after puromycin-based selection **(Figure 4D and S3, and Table S2)**. Most proteins that were significantly up-or down-regulated after palbociclib treatment followed by puromycin selection (P<0.05) did not reach statistical significance without selection **(Figure 4C and D)**. Cross-referencing our proteomics data with senescence protein databases showed that, in palbociclib-treated cells, differentially expressed proteins (P<0.05) overlapping CellAge ^23^ and SASP Atlas ^24^ databases were enriched ∼3.7-fold after puromycin selection **(Figure 4E and F)**. The most enriched protein in palbociclib-treated cells after selection was IGFBP2 **(Figure 4C and D)**, a protein previously reported as a SASP component ^24-26^ that is associated with ageing complications in humans ^27-29^. Therefore, we concluded that SenLect increased sensitivity for detecting upregulated senescence-associated proteins, as well as the metabolic and organellar rewiring that occurs in senescent cells.

**Figure 4.**
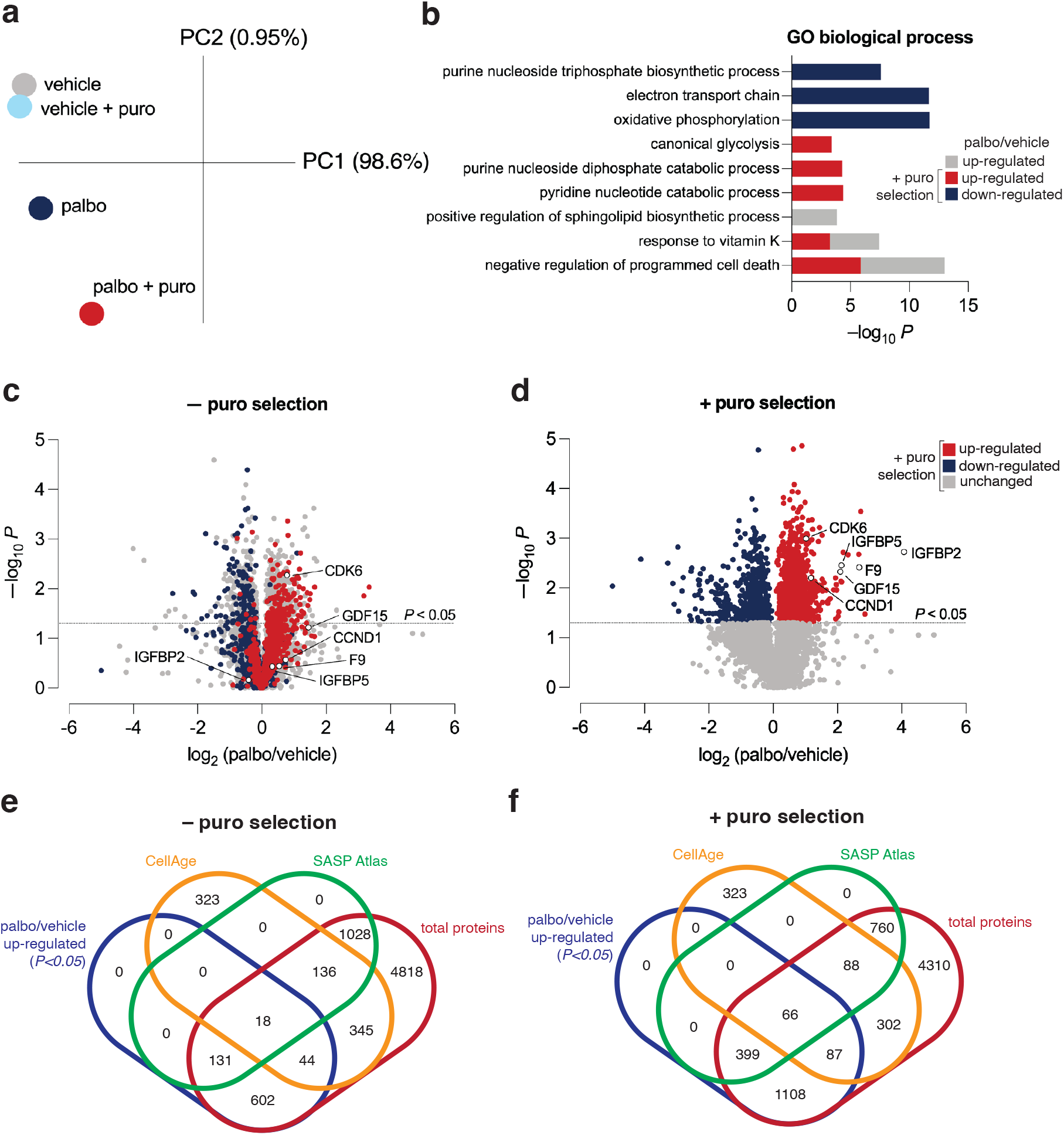
Coupling SenLect to global proteomics improves the resolution of senescence-related phenotypes after exposure to palbociclib. A. Principal component analysis of whole-cell proteomes of pSenLect-integrated HeLa cells following treatment with or without palbo after selection with or without puro. Summary of n=3 independent experiments. B. GO biological process analysis. Differentially expressed proteins (palbo/vehicle ≥2-fold up-or down-regulated; P<0.05) from whole-cell proteomes of pSenLect-integrated HeLa cells following treatment with or without palbo after selection with or without puro were used as search inputs. Selected GO terms are displayed; for comprehensive list refer to **Table S1**. Note that no GO terms were identified from the list of down-regulated proteins without puro selection. Summary of n=3 independent experiments. C and D. Volcano plots of whole-cell proteomes of pSenLect-integrated HeLa cells following treatment with or without palbo after selection with (D) or without puro (C). Log2 fold-change values >5 or <-5 were set to +5 or -5, respectively. n=3 independent experiments. See also **Table S2**. E and F. Venn diagrams showing overlap of palbo/vehicle ≥2-fold up-regulated proteins (P<0.05) with CellAge and SASP Atlas databases. Venn diagrams were produced using Venny ^49^. Summary of n=3 independent experiments.

## DISCUSSION

We recognised that to elucidate the molecular landscape of senescence, we must overcome the technical challenge of purifying senescent cells – which do not divide – away from proliferating neighbour cells. To this end, we developed a genetically encoded miR-146a promoter-based reporter we termed ‘SenLect’ that enabled mCherry fluorescence-based detection of senescent cells and their one-step enrichment via puromycin-mediated elimination of proliferative cells. This system is robust and can be used to enrich both primary and immortal cell cultures for senescent cells that arise from replicative senescence, UV irradiation, oxidative stress or chemotherapeutic agents. As proof-of-concept, using SenLect we demonstrated that removing proliferative cells to enrich for senescent cells significantly improved the detection of senescence-related signatures (e.g., SASP atlas, and CellAge) ^23,24^, which were otherwise hidden by the presence of proliferative cells, as revealed by whole-cell proteomics. This approach revealed the extent of proteome rewiring during senescence caused by cyclin-dependent kinase 4/6 inhibition, showing a shift in protein signatures from mitochondrial respiration and nucleotide synthesis toward glycolysis and nucleotide catabolism ^22^. It also revealed sub-proteome level rewiring of organelles like mitochondria, lysosomes and nucleolus, in addition to protein cohorts like kinases and core-essential proteins.

SenLect offers several advantages for purifying senescent cells compared to methods that isolate based on phenotypes such as large cell size (FSC), lipofuscin autofluorescence (Sudan Black-B analog GLF16), high senescence-associated β-galactosidase activity (C12FDG^+^) or plasma membrane receptor expression (TNFRSF10D/DCR2) using FACS or magnetic separation ^12-14,30-32^. All of these phenotypes can vary in a stimulus-dependent manner. Moreover, unlike the commonly used X-Gal approach ^33^, SenLect does not require cells to be fixed, meaning that live cells can be used for downstream experiments. It also provides an mCherry^+^ fluorescent readout of senescence, allowing this cell-state transition to be tracked in real time. Direct selection of senescent cells with SenLect bypasses the need for FACS. This is important because flow cytometry can lead to significant loss of senescent cells which are inherently fragile and also increases the risk of contamination. As such, SenLect is better suited for downstream experiments to be performed immediately after selection. SenLect is cost-effective and versatile: it provides consistent senescent cell enrichment for different stimuli in both primary and immortalised cell lines. It is likely to be broadly useful and highly compatible with diverse downstream applications, including RNA-seq, phospho-proteomics, CRISPR-Cas9 screens, and functional assays etc. We also suggest that miR-146a-promoter systems may be a better alternative to the commonly used p16- and p21-promoter systems for detecting senescent cells in living organisms ^34-37^. This is because some cell types and cancers naturally show high levels of p16 or p21, making those markers harder to interpret in such situations. Senescent cells are generally rare *in vivo* and make up <10% in most aged tissues of mice and humans, except the endocrine pancreas where they make up ∼35% of cells ^38-40^. SenLect can enrich these rare populations of senescent cells in culture. Therefore, developing a SenLect mouse would allow the study of senescence across different cell types by generating primary cell cultures from various tissues and then selecting for their constituent senescent cells *ex vivo*.

In summary, SenLect offers a practical way to separate live senescent cells from dividing cells in culture, making it useful for a variety of downstream experiments. It permits a more detailed view of the molecular features of senescent cells, which could help identify new, more sensitive biomarkers of senescence relevant to human health.

## Limitations

The SenLect system has some technical limitations that may affect its use in certain situations. Since it is genetically encoded, it needs to be introduced into cells using lentiviral transduction. This can be challenging for cell types that are hard to transduce, and using a high virus dose may cause the system to become ‘leaky’. It may be possible to manage this by selecting low-expressing clones, using a minimal promoter sequence or using CRISPR-Cas9 to knock-in mCherry-P2A-PuroR directly downstream of the endogenous miR-146a promoter. SenLect is most useful when only a small-to medium fraction of cells in the culture are senescent. Finally, the miR-146a promoter is only active in senescent cells that produce the SASP, which may limit its usefulness for isolating senescent cells that do not have a SASP ^41^.

## MATERIALS AND METHODS

### Plasmid construction

pSenLect (VB240218-1391uev), pSenLect_+Control (VB240220-1398dkj), and pSenLect_-Control (VB240225-1125wmh) were synthesised in the pLV[Exp] lentivector backbone by VectorBuilder.

The *Homo Sapiens* miR146 promoter sequence (1.5 kb; -1.5 kb from the 5’ region of exon 1) was used:

5’_CTATATAGCTATAGCTCAATAAGGCTATTATATACTTATTTTATTTTATTATTATTTTTTAGAGACAGAGTCTCGCTTTGTCACCCAGCCTGGAGTGCAGTTGTGCAATCTTAGCTCACTGCAACCTCTGCCTCCTGGGTTCAAGTGATTCTTCTGCCTCAGCCTCCCAAGTAGCTGGGACTACAGGTGTGCGCCACCATGCCTGGCGAATTTTTGTATTTTTGTTAGAGACGGGGTTTCACCATGTTAGCCAAGTGAAACTTGAACTCCTGACCTCAGGTGATCTGCCTGCCTCAGCCTCCCAAAGTTCCGAGATTACAGGTGTGAGTCACCACATCTCGCCTATATACATATTTAAAAAACACTAGCATGGTCTCTATGCTTATGGAAAACAGTCTTGTTGGAGAGATAATTTTAATCAAATAATTATACAAAATTATCTAACAATTACAAATTACGGTGAAGGAAAGAATCCCATATTATGAAAGAGACTAGCTGGAGGCTGATGGTGAGGGAGTGGTAGGTAAACATTCAAGTGCTCTTACAGAGATCCCCAAGTAACAGTGGCTTAAATAAGGTAAATGTTCCATTCTTGCTGATCTATCAGTGCAGGTGTAAGCAGTCTAGGGTGTATAAGGAAGCCTCACACTTTTGGGGCTCTTCTGGCATGCTGCTCTGCTGTTCCCTGGGGGGCTGCACTCATTTGCATGGTCCACAGTGGCTCTCTACCATGTCTACAGTCCAAGCCTGAGAAGAGGGAGGGAAACAGTAAAAAACTCTCCACTCCTGCTCATATCCAGCACTTTGTCACACCAGGTGAGGAAACAAAGTCAGATGTCCATGTGTTCAGATACCACTTAAGGGTTCTATTAGTATAGAAGGAGGAGAGAATATATATTGGGCAAACAATCAGCAATCTCTGCCATAGATGGCCTCTCTGAGGAAGTGACATTGAAAGCAAGAACATCAGGATAAGTTGATGTTAAAGAGAGGAACGAGTTTTCCAGGCCAGAGGGATGGCATATGGAAGGGTCATGAGGCAGGAAAGGCCAGCTACCATGTTTAATGATGCTTAATGATGTGTGTGTCTACCATACACATCCCCTACAGATTAGTTTTTGTTTTGACAGGGTCTCTCTCTGTGGCCCAGACTGGAGTGCAGTGGTGCAATCATAGCTCACTGCAACCTCCAATTCCCAGGCTCAAGCGATCCTCCCACCACAGGCCATCATGCATGGCTCATTTTTTATTTTTAGTAGAGACAAATTCTCCATGTTGCCCAGGCTAGTCCTGAACTCCTGGGCTCAAGAGATCCACCCACATCAGCCTTCCAGACTGCTGGCCTGGTCTCCTCCAGATGTTTATAACTCATGAGTGCCAGGACTAGACCTGGTACTAGGAAGCAGCTGCATTGGATTTACCAGGCTTTTCACTCTTGTATTTTACAGGGCTGGGACAGGCCTGGACTGCAAGGAGGGGTCTTTGCACCATCTCTGAAAA_3’

### Cell culture

HUVEC cells (cc-2517, Lonza) were maintained in Human Large Vessel Endothelial Cell Basal Medium (M200500, ThermoFisher) supplemented with Large Vessel Endothelial Supplement (LVES) (A1460801, ThermoFisher) at 37°C and 5% CO_2_. Cells between passages 4 and 15 were used for these experiments. HeLa cells were maintained in DMEM (11965092, ThermoFisher) supplemented with 10% FCS (SAH32202-20-1, CellSera). Cells were routinely screened for mycoplasma contamination using MycoAlert (LT07-318, Lonza).

### Lentiviral Transduction

For each construct, HEK293T cells were seeded in two wells of a six-well plate at a concentration of 5 × 10^5^ cells per well and incubated overnight. The following day, cells were transfected with 5 µg of the pSenLect lentivector (or its controls), 4 µg of psPAX2 (Addgene plasmid #12260, a gift from Didier Trono), and 4 µg of pCMV-VSV-G (Addgene plasmid #8454, a gift from Bob Weinberg) using Lipofectamine 3000, according to the manufacturer’s protocol, and incubated for 48 hours.

HUVECs (passages 5-7), used as target cells, were seeded at 1.5 × 10^5^ cells per well in two wells of a six-well plate one day prior to transduction. On the day of transduction, the viral supernatant from the transfected HEK293T cells was harvested, filtered, and supplemented with polybrene to a final concentration of 8 µg/mL before being added to the HUVECs. The following day, the viral medium was replaced with complete media. After 72 hours, fluorescence microscopy revealed TagBFP^+^ fluorescence in the transduced cells, confirming successful expression. Blasticidin S was then added at a concentration of 4 µg/mL to select for transduced cells, and cells were maintained under selection in 2 µg/mL blasticidin S (except when being used in experiments).

### Generation of stable SenLect monoclonal HeLa cells

HeLa cells were transduced similar to that described above, except 3 × 10^5^ cells per well were plated. After blasticidin S selection, single TagBFP^+^ cells were sorted into a 96-well plate using a BD FACS Melody cell sorter. Each well contained 200 µL of DMEM supplemented with 20% FCS and 1% penicillin/streptomycin. After three-weeks, single clones were expanded, and duplicate plates prepared to test their mCherry^+^ responsivity to senescence induction by doxorubicin which was measured by flow cytometry (BD Fortessa). The clone with the highest mCherry expression under senescence conditions was used for subsequent experiments.

### Senescence induction

Senescence was induced in HUVECs by exposure to UV (312 nm) for 20 seconds on three consecutive days while cells were in DPBS. Alternatively, cells were treated with 50 µM H_2_O_2_ (HA154-500M, Chemsupply) or 200 nM doxorubicin (100-0558, Stem cell technologies) for 72 hours. In HeLa cells, senescence was induced by treatment with 7.5 μM palbociclib (HY-50767A, Med Chem Express) for 72 hours, or 200 nM doxorubicin for 24 hours followed by 48 hours washout. Corresponding vehicle solutions were used as a negative control. At the end of the experiment, senescence was assessed by cell size, cell granularirty, senescence-associated β-galactosidase activity, mCherry^+^ fluorescence, and cell cycle status.

### Selection of senescent cells

After 72 hours of senescence induction, the cell culture medium was replaced with fresh medium with or without 0.5 µg/mL or 1 µg/mL puromycin (P8833, Sigma) for 72 hours for HUVECs or HeLa cells, respectively. This eliminated non-senescent cells that did not exhibit puromycin resistance.

### Senescence-associated β-galactosidase activity and cell size

Senescence-associated β-galactosidase activity was measured using C12FDG fluorescence (ab273642, Abcam). On the final day of the experiment, cells were incubated with 100 nM bafilomycin (S1413, Sapphire Bioscience) for 1 hour in fresh culture medium. Following this, 2 mM C12FDG was added to the medium to achieve a final concentration of 33 µM; cells were then incubated for 1 hour or 2 hours for HUVECs and HeLa cells, respectively. After incubation, the cells were harvested, resuspended in DPBS, and analysed by flow cytometry (BD Fortessa) ^42^. Flow cytometry was also used to analyse mCherry fluorescence, cell size (FSC), and granularity (SSC). Cells transduced with pSenLect_+Control and pSenLect_-Control were used as controls for establishing mCherry positive and negative gates.

### Cell cycle analysis

After flow cytometry for C12FDG, the remaining unanalysed cells were centrifuged to discard the DPBS. To the cells, 1 mL of cold 70% ethanol was added drop-wise while vortexing, then stored at 4°C. On the day of staining, fixed cells were centrifuged at 2,700 rpm for 5 min to remove ethanol and incubated with 50-100 µL of 100 µg/mL RNase A (EN0531, ThermoFisher) for 5 min at room temperature. Following this, 200-400 µL of 50 µg/mL propidium iodide (P3566, Thermofisher) was added to each sample, vortexed, then incubated in the dark at 37°C for 45 min. The cell cycle distribution was evaluated using flow cytometry (BD Fortessa).

### Mass spectrometric (MS) Analysis

Cells were harvested and prepared using an EasyPep™ MS Sample Prep Kit (A45735, ThermoFisher) according to the manufacturer’s instructions. After peptide cleanup, samples were dried in a vacuum centrifuge and reconstituted in LC loading buffer (2% acetonitrile, 0.1% trifluoracetic acid) at a concentration of 200 ng/µL. Using an Acquity UPLC M-Class System (Waters Corporation), an injection volume of 1 µL was loaded onto a Waters nanoEase M/Z Symmetry C18 trap column (20 mm x 180 µm, 5 µm) for 5 minutes at 10 µL/min using LC loading buffer before transferring to an IonOpticks Aurora Ultimate CSI UHPLC column (25 cm x 75 µm, 1.7 µm) for analytical separation. The following chromatography gradient was used: 40 min linear separation from 3-40% mobile phase B (0.1% formic acid in acetonitrile), followed by a 10 min wash at 95% mobile phase B and re-equilibration at starting conditions for 15 min. Mobile phase A consisted of 0.1% formic acid in water.

All MS data was collected using a timsTOF Pro2 (Bruker) in positive ionisation mode with the dia-PASEF-long gradient default method. All samples were acquired in a random order.

### Organelle and protein class annotation of proteomics data

We curated a master list of proteins that belong to various cellular organelle compartments and protein classes which have been experimentally validated ^43-48^ or are available from online databases:

https://esbl.nhlbi.nih.gov/Databases/KSBP2/Targets/Lists/DUBs/

http://www.kinhub.org/kinases.html

### Data analysis

Flow cytometry data were analysed using FlowJo™ version 10.8 (BD Life Sciences). Statistical analyses were conducted with GraphPad Prism version 10. Specific sample sizes and statistical tests for each experiment are described in the figure legends. Proteomics raw MS data were analysed against the human proteome FASTA database (UniProt) via Spectronaut version 20 (Biognosys) directDIA using default settings without cross-run normalisation. MetaboAnalyst version 6.0 (metaboanalyst.ca), Microsoft Excel and GraphPad Prism were used for further data visualisation and statistical analyses. A *p-value* of ≤ 0.05 was considered statistically significant.

## Supporting information

Tabl1 S1

## ABBREVIATIONS

BlastR: blasticidin S resistance/blasticidin S deaminase
C12FDG: 5-dodecanoylaminofluorescein di-β-D-galactopyranoside
FACS: fluorescence-activated cell sorting
FSC: forward scatter
HUVECs: Human Umbilical Vein Endothelial Cells
MS: Mass spectrometric
PuroR: puromycin resistance/puromycin N-acetyltransferase
SASP: senescence-associated secretory phenotype
SenLect: senescence selection
SSC: side scatter.

## ACKNOWLEDGMENTS

This investigation was supported by Lysosomal Health in Ageing at SAHMRI. JMC is supported by an EMCR Fellowship from The Hospital Research Foundation Group (2022-CF-EMCR-007). We thank all members of Lysosomal Health in Ageing (SAHMRI) for helpful insights and discussions. We thank Alexis Martin and Dr Sanjna Singh (SAHMRI) for training H.T. in some experimental techniques; Dr Marten Snel from the Proteomics, Metabolomics and MS-Imaging Facility (SAHMRI) for advice in the interpretation of proteomics data; and Dr Randall Grose and Sophie Thomson for flow cytometry support. Flow cytometry analysis was performed at the Adelaide Health and Biomedical Precinct Cytometry Facility (SAHMRI), which is generously supported by the Detmold Group, the McMahon Family, the Australian Cancer Research Foundation, the Cancer Council, and the Australian Government through the Zero Childhood Cancer Program.

---

**Figure S1.**
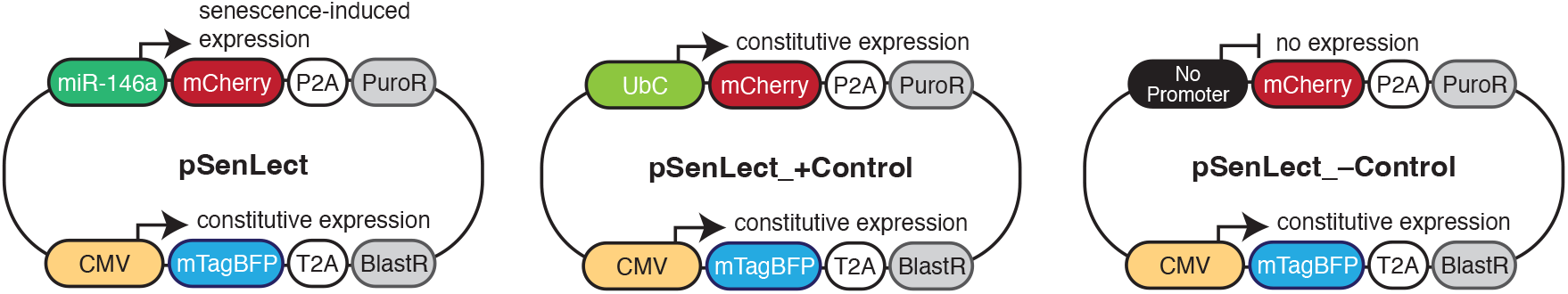
**Comparison of pSenLect and its positive and negative controls plasmids that are useful for establishing flow cytometry gating**.

**Figure S2.**
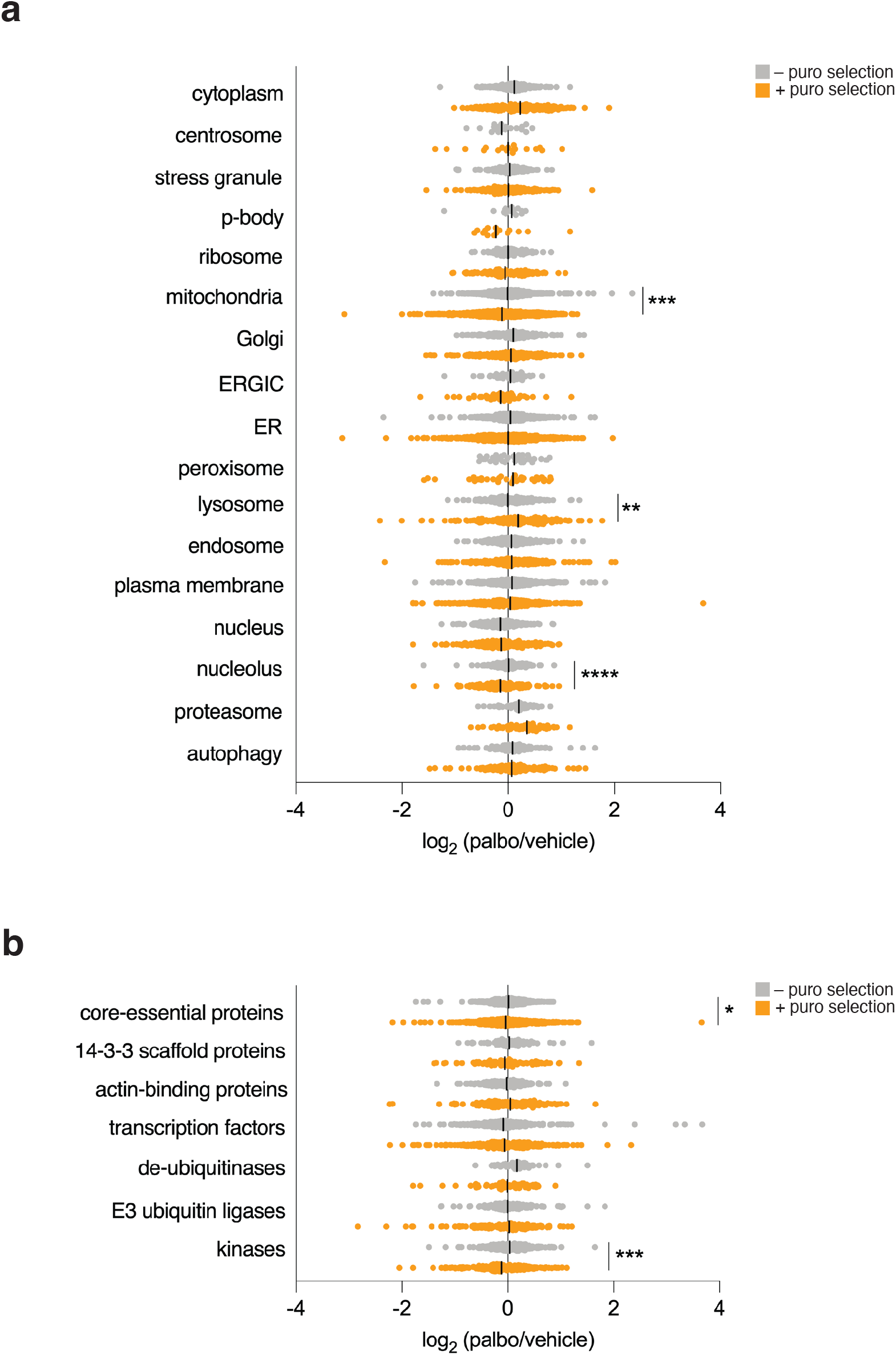
**Sub-proteomes of organellar comparments (A) and protein cohorts (B) from whole-cell proteomes of pSenLect-integrated HeLa cells following treatment with or without palbo after selection with or without puro. Summary of n=3 independent experiments**.

**Figure S3.**
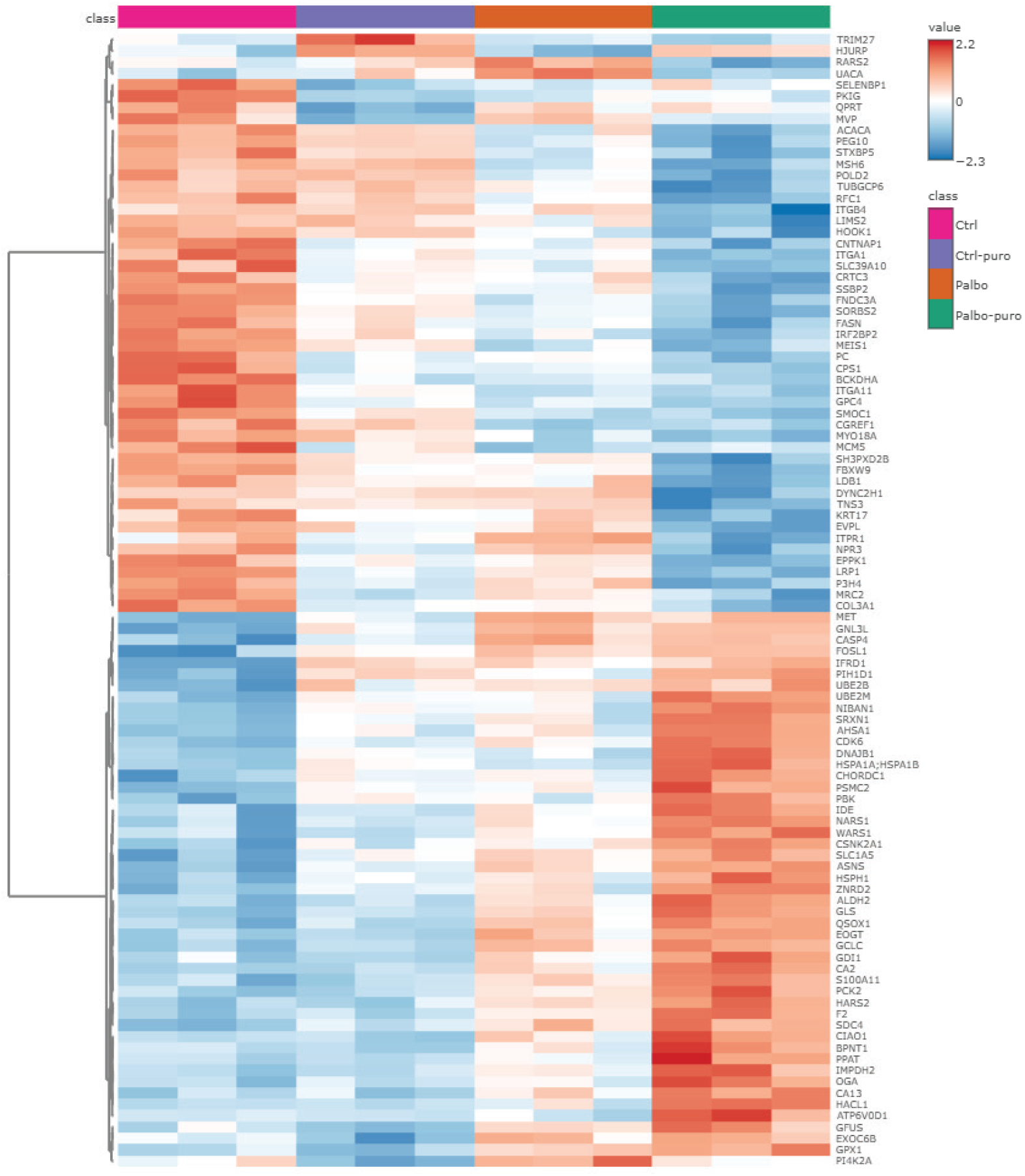
**Heatmap of top and bottom 50 differentially expressed proteins (palbo/vehicle; P<0.05) from whole-cell proteomes of pSenLect-integrated HeLa cells following treatment with or without palbo after selection with or without puro. Summary of n=3 independent experiments**.

**Table S1. GO Biological Process Complete and Reactome analyses of differentially expressed proteins from whole-cell proteomes of pSenLect-integrated HeLa cells following treatment with or without palbo after selection with or without puro. Proteins with fold-change (palbo/vehicle) ≥2 up-or down-regulated (P<0.05) were used as search inputs**.

**Table S2. Raw data for used to prepare Volcano plots in Figure 4C and D**.

